# MMP release following cartilage injury leads to collagen loss in intact tissue – a computational study

**DOI:** 10.1101/2025.06.10.658780

**Authors:** Moustafa Hamada, Atte S.A. Eskelinen, Joonas P. Kosonen, Cristina Florea, Alan J. Grodzinsky, Petri Tanska, Rami K. Korhonen

## Abstract

Damage of collagen fibril network in articular cartilage plays a key role in post-traumatic osteoarthritis but the main underlying mechanobiological mechanisms of fibrils degeneration early after injury are not fully understood. This study explores the hypothesis that injurious loading leads to cellular damage that triggers the release of matrix metalloproteinases (MMPs), resulting in loss of collagen content in cartilage. To investigate this, we developed a computational mechano-signaling model simulating spatial collagen loss in bovine cartilage. In the model, the injurious loading causes excessive shear strains in tissue matrix, leading to cell damage and subsequent release of MMPs. The model was compared to *ex vivo* cartilage explant experiments over 12 days post-injury where collagen content was assessed via Fourier-transform infrared microspectroscpy. By day 12, the simulated collagen loss aligned with our experimental findings along most of tissue depth (∼30% bulk average loss in the model vs. ∼35% in the experiment). The results suggest that injury-related cell damage and the downstream MMP activity could partly explain the depth-wise collagen content loss in early days after *ex vivo* cartilage injury. Ultimately, combining the current approach with joint-level computational models could enhance the prediction of the onset and progression of cartilage degeneration.

**Author Summary:** Knee injuries can initiate an irreversible degeneration of articular cartilage which can later lead to the development of post-traumatic osteoarthritis over the years. Currently, there is no cure or effective intervention to repair degenerated cartilage or halt disease progression. Yet, opportunities for intervention depend on a thorough understanding of mechanisms that govern the very early changes in cartilage tissue composition such as the collagen fibrils. In this work, we developed a finite element computational modeling framework to simulate cartilage following injurious loading, focusing on the early loss of collagen content. The model aims to capture the role of injury-induced cellular damage in elevating proteolytic activity within the tissue, which could rapidly degrade collagen fibrils and result in significant collagen loss. This model framework offers a tool for studying the degradation of the collagen fibril network, testing hypotheses involving cell-driven mechano-signalling pathways and evaluating potential treatment interventions aimed at preventing collagen loss.

## 1. Introduction

Post-traumatic osteoarthritis (PTOA) is a subtype of osteoarthritis (OA) that can develop in response to joint injuries, such as anterior cruciate ligament injury, meniscus tears, or intra-articular fracture[1]. While timely clinical intervention after injury can restore joint function, altered biomechanics, pro-longed inflammatory response, and compositional changes in articular cartilage tissue contribute to the long-term risk of developing PTOA[1–3]. PTOA can have devastating consequences, particularly for active and athletic individuals[3]. Symptoms often emerge years after the initial injury, presenting as joint instability, limited range of motion and narrowing of joint space[2,3]. In advanced stages, the disease may result in joint pain or complete disability, eventually necessitating joint replacement[2]. A key factor in advancing interventions for this disease is a detailed understanding of cartilage mechanobiology and the elucidation of mechanisms that govern cell-tissue interactions in the early stages following injury. Multiple experimental and computational works suggest that cartilage degeneration is initiated by cellular cues that lead to cartilage matrix loss[4–7]. However, the link between early-stage cell damage after injury and the degradation of cartilage collagen fibrils, as well as the spatial and temporal progression of this degradation within the tissue remains ambiguous.

In response to injury, cell damage in cartilage can be represented by mitochondrial dysfunction, oxidative stress, and increased catabolic activity[7–9]. On the tissue level, injured cartilage can be characterized by aggrecan loss and collagen fibril network degradation[10–13]. This cascade of cellular and tissue-level responses can be interconnected through mechano-signaling pathways. *Ex vivo* investigations have provided strong evidence that injury-induced cell damage, the imbalance between catabolic and anabolic chondrocyte activities, and increased production of matrix metalloproteinases (MMPs) and aggrecanases could later lead to enzymatic degradation of collagen fibril network and aggrecan loss, respectively[7,9,14–16]. Specifically, elevated mechanical shear strains within the tissue can be co-localized with cellular damage such as mitochondrial dysfunction, production of reactive oxygen species, and cell apoptosis[4,9,17,18]. The high cellular strains can activate mechano-signaling pathways through cell surface receptors, such as integrins, low-density lipoprotein and primary cilia[19–21]. This may culminate in the upregulation of MMPs which can be detected with immunohistochemical staining[22]. The secreted MMPs such as MMP-1, MMP-3, and MMP-13[16,23] can cleave intact and mechanically ruptured collagen fibrils by binding to specific sites within the collagen triple helix structure, generating three-quarter and one-quarter length fragments[24]. These collagen fragments can then undergo further degradation by other MMPs such as MMP-2, and MMP-9[25]. Ultimately, this cascade can lead to collagen content loss[13,16,20,25,26] which is observable along tissue depth and near chondral lesions as early as 12 days from injurious loading[22]. On the other hand, these catabolic protease activities can also be downregulated with several inhibitory or anabolic factors such as the increased expression of tissue inhibitors of metalloproteinases (TIMPs)[20] or blocking of MMPs from binding to collagen fibrils cleavage sites by aggrecan molecules[27].

Mechanobiological models offer a promising avenue for investigating the driving mechanisms behind early PTOA-related compositional changes, which are challenging to attain solely by experiments[26,28–30]. When it comes to collagen damage, a major focus of current modeling studies is typically on collagen mechanics such as collagen network softening/damage driven by excessive tensile stresses in the cartilage matrix or tensile strains in the individual collagen fibrils[31,32]. On the other hand, several computational studies have addressed cellular activity and biochemical signaling that could lead to the loss of collagen fibrils[8,33–35]. However, these models overlooked the possible damage localization and its progression over time after injurious loading[26,28,34].

In this work, we developed a cell-driven mechano-signaling modeling framework to simulate depth-wise loss of collagen content observed in cartilage explants exposed to injurious loading[22]. We hypothesized that injurious loading would result in excessive shear strains triggering cell damage. This can lead to an increase in MMP production, and finally enzymatic cleavage of collagen fibrils[4,18]. The model included basal collagen loss in the control intact cartilage explants by calibration of the experimentally observed depth-wise loss of collagen content at day 12 in control free-swelling conditions[22].

It has been suggested that aggrecan molecules may confer a protective effect by blocking MMP access to collagen fibrils, thereby slowing down collagen loss[27]. Our model incorporated this modulatory role of aggrecan to the MMP-mediated collagen loss. Furthermore, we simulated aggrecan loss as a consequence of aggrecanse-driven enzymatic degradation, under the assumption that a reduction in aggrecan content would affect this protective effect and facilitate increased collagen loss[27,28]. To verify model results, the simulations were compared against measured depth-wise collagen content in injuriously loaded young bovine cartilage explants on the day of the injury and 12 days after the injury[22].

## 2. Methods

### 2.1 Experiment design

This study utilizes a subset of data from previous experiments[22]. Briefly, cylindrical cartilage explants (*n* = 63, diameter = 3 mm, thickness = 1 mm) were harvested from four different regions of the patellofemoral grooves of two-week-old bovines (*N* = 8)[36,37]. A subset of explants was subjected to injurious loading (50% strain, 100%/s strain rate) to be assessed for collagen content on the day of injury (day 0, *n* = 18), and after 12 days of incubation in culture medium (*n* = 6). The remaining explants were kept as a free-swelling control group (day 0: *n* = 18 and day 12: *n* = 21).

### 2.2 Experimental collagen content assessment

Fourier-transform infrared microspectroscopy (FTIR, 5.5 × 5.5 µm^2^ pixel size, 4 cm^-1^ spectral resolution) was used to assess the spatial collagen content distribution[22]. Three 5-µm-thick unstained tissue sections were prepared for the measurements per explant. The area under Amide I spectral region (wavenumber range: 1580 cm^-1^ to 1720 cm^-1^) was calculated pixel by pixel from each section per explant and used as an estimate for the collagen content (Fig 1a)[38,39]. Next, two full-depth and 200 µm wide regions of interest (ROI) were defined in each histological section. Collagen content was extracted from the ROIs, averaged perpendicular to depth, and normalized depth-dependently (0% = surface, 100% = bottom) to obtain depth-wise collagen content for each explant. The results are presented for each group as the mean depth-wise collagen content with ±95% confidence intervals (Fig 1b) [22]. To assess the average decrease in collagen content in whole tissue, the mean value is calculated from the entire ROI for each explant. Analysis of the aggrecan content of the same explants were done and results reported earlier in a previous study[36].

**Fig. 1.**
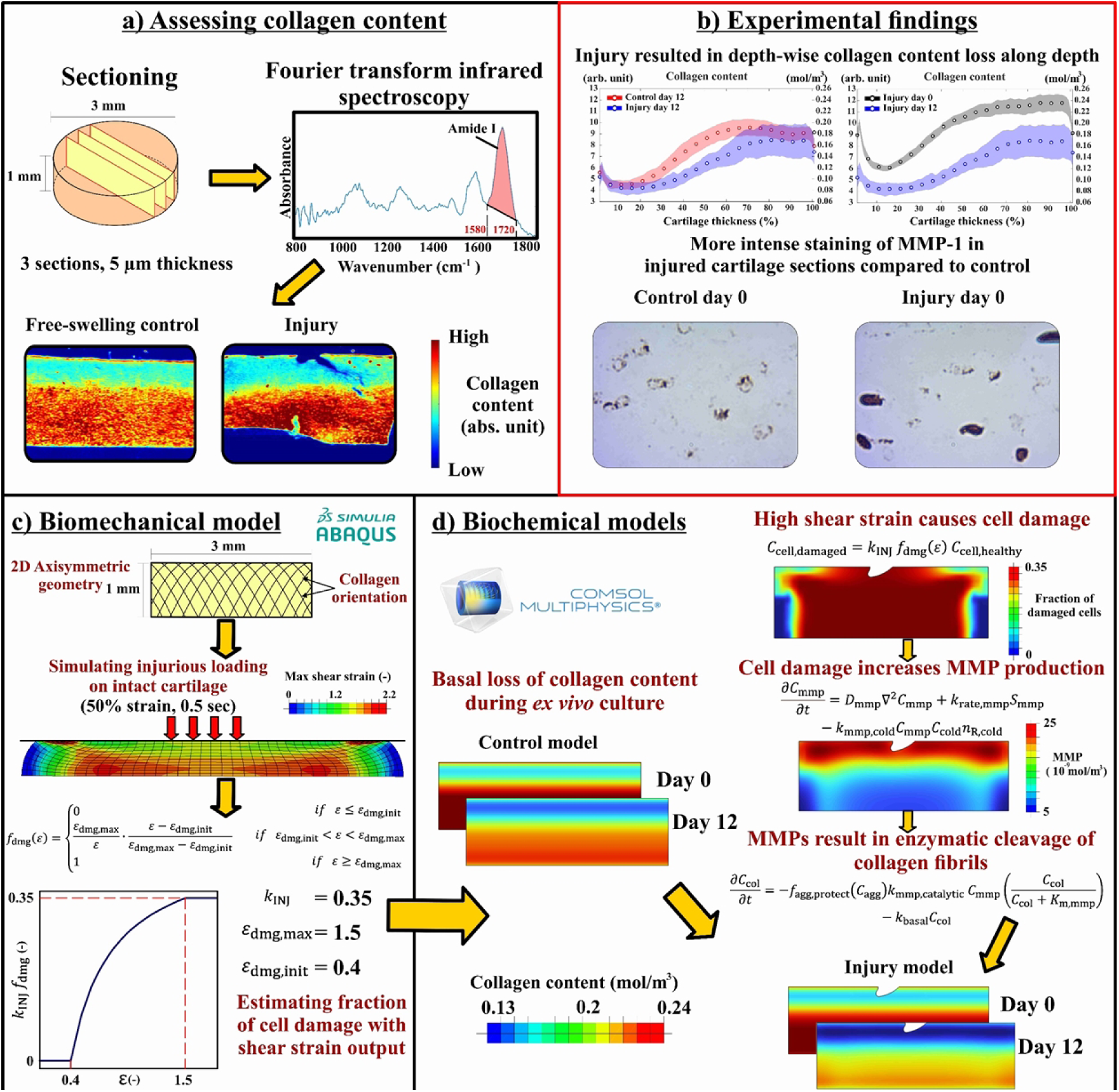
Study workflow. **a)** Collagen content was assessed using Fourier-transform infrared microspectroscopy from histological sections of control free-swelling and injuriously loaded cartilage explants. **b)** Key experimental findings showed that injurious loading can result in depth-wise collagen content loss 12 days after injury when compared to control group and injuriously loaded group on day 0. In addition, immunohistochemical analysis showed more intense staining of MMP-1 in injured cartilage compared to intact control. **c)** Injurious loading was simulated with finite element model to obtain maximum shear strain distribution and estimate fraction of damaged cells. **d)** Collagen content loss was modeled over 12 days following injurious loading with reaction–diffusion equations.

### 2.3 Biomechanical computational model

A 2D finite element (FE) model was created (Fig 1c) using ABAQUS software (v. 2023, Dassault Systèmes, Providence, RI, USA) to simulate injurious loading of an intact cartilage explant under unconfined compression. The 2D geometry of intact cartilage was created as in the experiment (diameter = 3 mm, thickness = 1 mm).

Cartilage explant was meshed using 456 linear quadrilateral elements of type CPE4P. Prior to model implementation, different mesh densities were tested to ensure the model results were mesh-independent[40]. The cartilage was modeled as a fibril-reinforced porohyperelastic material with swelling, and the material parameters optimized based on experimental data from similar young bovine cartilage explants (Table 1)[37,41].

**Table 1.** Main material parameters for the biomechanical model. *z* indicates normalized distance from the cartilage surface.

| Material parameter | Value | Description | Reference |
| --- | --- | --- | --- |
| $E_f$ (MPa) | 20.0 | Fibril network modulus | [37] |
| $C$ (-) | 3.009 | Relative density between primary and secondary fibrils | [32] |
| $E_m$ (MPa) | 0.16 | Non-fibrillar matrix modulus | [37] |
| $\nu_{nf}$ (-) | 0.4 | Poisson's ratio of the non-fibrillar matrix | [57] |
| $k$<br>($10^{-15} \text{ m}^4/(\text{N}\cdot\text{s})$) | 1.3 | Hydraulic permeability | [37] |
| Distributions | Value | Description | Reference |
| $n_{f,0}$ | $0.85 - 0.1z$ | Initial depth-wise fluid fraction | [37] |
| $c_{col,0}$ | $-187.1z^6 + 328.6z^5 + 22.9z^4 - 363.4z^3 + 247.8z^2 - 47.9z + 8.782$ | Initial depth-wise collagen fraction | [22] |
| $c_{FCD,0}$ (mEq · ml<sup>-1</sup>) | $-4.4z^6 + 15.2z^5 - 21.0z^4 + 14.9z^3 - 5.8z^2 + 1.1z + 0.03$ | Initial depth-wise fixed charge density | [37] |

The collagen fibrils network was modeled as a combination of large primary fibrils and smaller secondary fibrils[32]. The primary fibrils orientation constructed based on earlier polarized light microscopy measurements of fibril orientation of our young bovine cartilage samples[37,42], where fibril orientation to the surface is ∼10^ο^ and linearly reaches ∼55^ο^ at the bottom of the tissue (Figure 1c)[37]. On the other hand, the secondary fibrils were modeled to define randomly oriented fibrils[32]. The initial depth-wise collagen fraction distribution (*c*_col,0_) was implemented in the model based on the mean depth-wise collagen content in intact cartilage explants at day 0 (Table 1)[22].

The collagen fibrils were modeled as linear elastic where the stress on the fibril (*σ*_f_) is governed by:

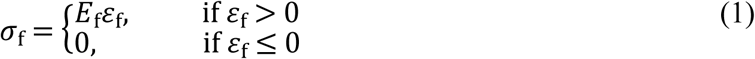

where *E*_f_ is the fibril network modulus and *ɛ*_f_is fibril strain. When considering the relative density between primary and secondary fibrils (C), the stresses in the primary (*σ*_f,p_) and secondary fibrils (*σ*_f,s_) can be given by:

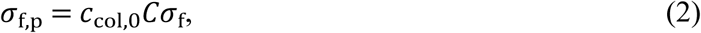

and

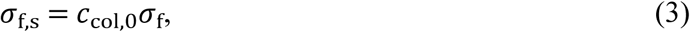

Similar to collagen fibrils distribution, the depth-wise fixed charge density (FCD) distribution of proteoglycans was implemented using optical density measurements from the same cartilage explants (characterized in[37]) and converted into FCD values as done previously[43]. The fluid fraction was obtained from literature[43].

#### 2.3.1 Boundary and loading conditions

The bottom surface of the explant was fixed in the axial direction, while a single node at the intersection point between the center line of the explant and the bottom surface was fixed in the lateral direction to fix the model in place but allow lateral expansion of the tissue simultaneously (Fig 1c). The contact between the tissue’s top surface and the compressing plate was modeled as frictionless surface-to-surface interaction. Free fluid flow was allowed from the lateral edges (pore pressure was set to 0). Finally, similarly as in the experiments, a single load-unload cycle with a 50% strain magnitude (strain rate 100%/s, 50% strain reached at 0.5 s) was applied on the top surface of the explant to simulate the injurious loading.

### 2.4 Biochemical computational model

A 2D geometry of cartilage explant was created using COMSOL Multiphysics (version 6.1, MA, USA) with same dimensions as in the biomechanical model. We did not model lesion formation or crack propagation in the mechanical model, but lesion geometry was added to the injury model geometry (Fig 1d). The lesion geometry was based on microscopy measurements of a representative sample after injurious loading[22,36,37]. It was created to visually assess the localization of collagen loss around this region after injury.

#### 2.4.1 Governing equations

To simulate cell damage and subsequent collagen loss over 12 days, the maximum shear strain distribution at 50% of compressive strain was obtained from the biomechanical model by calculating the maximum difference between the principal components (*ɛ*_*p*,1_, *ɛ*_*p*,2_, *ɛ*_*p*,3_) of the Green–Lagrangian strain tensor[29]:

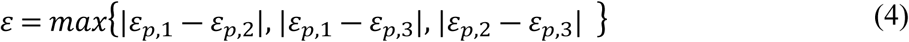

The maximum shear strain values were used as they have been linked with chondrocyte damage in deformed tissue regions[37]. A stepwise non-linear function[18] *f*_dmg_(*ɛ*) was used to represent cell damage dependency on the amount of shear strain (Fig 1c):

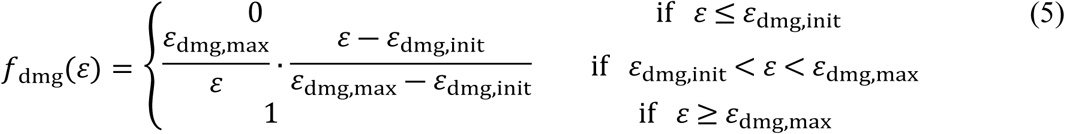

in which we assumed that cells start to turn damaged when the maximum shear strain value exceeds 40% (ε_dmg,init_ = 0.4) and after that, the amount of damaged cells increases non-linearly as a function of strain until the maximum fraction of damaged cells is reached at the maximum shear strain value of 150% (ε_dmg,max_ = 1.5). These threshold values are based on previous computational studies using fibril-reinforced poroelastic material models and experimental data using excessive levels of strain (indentation test)[9,37]. Subsequently, the concentration of damaged cells *C*_cell, damaged_ was defined as a fraction of the concentration of healthy cells *C*_cell, healthy_:

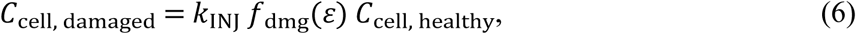

where *k*_INJ_ = 0.35 is a multiplier to limit the maximum fraction of cell damage in the model to 35% from healthy cells. This estimate of cell damage is adopted from previous cell apoptosis measurements in injured young bovine cartilage[23].

The temporal and spatial changes of the concentrations of the cartilage structural components, MMPs and aggrecanases post-injury were modeled with reaction-diffusion equations as[28]:

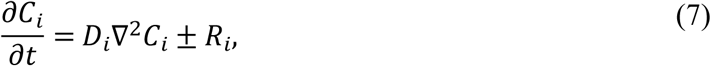

where *C*_*i*_indicates the concentration of a species *i*. Species considered in the model are intact collagen (col), degraded collagen (cold), intact aggrecan (agg), degraded aggrecan (aggd), MMPs (mmp), and aggrecanase (aga). *D*_*i*_is the effective diffusivity of the species *i* while *R*_*i*_ denotes a variable that encompasses the rates of production or degradation of the species *i*[28]. We only considered the net effect of proteases and the different types of MMPs (such as MMP-1, and MMP-3) or aggrecanases (such as ADAMTS-4 and ADAMTS-5) were not considered for model simplicity.

As the activation and secretion of MMPs and aggrecanases after injury is not instantaneous, time-dependent stimulus variables *S*_mmp_ and *S*_aga_ were introduced to represent the time delay of MMP and aggrecanase activations, respectively[28]:

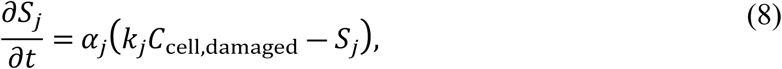

where *α*_*j*_ corresponds to the rate constant for the net stimulus of protease *j* (*j* = mmp or aga), and *k*_*j*_ is a constant for protease released from damaged cells. The subsequent change in MMP production over time was governed by:

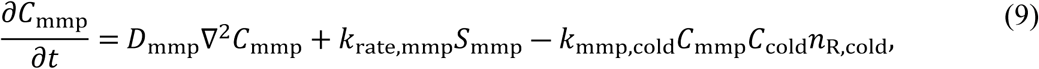

where *k*_rate,mmp_ is the rate constant for generating MMPs based on damaged cell-driven stimulus, *k*_mmp,cold_is a rate constant that consider the decrease of active MMPs to cleave intact collagen fibrils due to binding to degraded collagen[26,28] and *n*_R,cold_ represents the number of MMP binding sites on degraded collagen[26]. The model assumes that the release of MMPs occurs at the same rate over the 12-day simulation. The change in aggrecanases production follows the same approach and it is described in more detail in equation (S.E2) in supplementary material.

Collagen fibrils possess multiple protection mechanisms that help in hindering collagen degradation. One suggested mechanism is that aggrecan molecules can bind to type III and IV collagen on the surface of type II collagen fibrils[27]. This can block MMPs molecules from reaching binding sites on the fibrils and mitigating collagen degradation. This effect was incorporated in the model with a function *f*_agg,protect_(*C*_agg_) that defines how local aggrecan concentration limits the tendency of MMPs to cleave collagen fibrils[28] and can be described by:

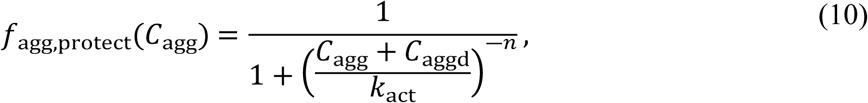

where *C*_agg_and *C*_aggd_are the concentrations of intact and degraded aggrecan respectively, *k*_act_ defines aggrecan concentration at half-maximal MMP activity, and *n* is the Hill coefficient of MMP activity[28]. Changes in intact and degraded aggrecan concentrations are defined in equations (S.E3) and (S.E4), respectively in the supplementary.

The *ex vivo* incubation of cartilage explants in culture media can maintain tissue homeostasis and structure intact for weeks. Yet, an inherent basal degradation of cartilage components might be inevitable with small level of aggrecan loss and cell death reported earlier in freshly cut explants[28,35,36,44]. We accounted in the model for the potential basal collagen loss with a rate constant *k*_basal_. The model was calibrated for this basal depth-dependent loss by adjusting *k*_basal_ rate constant till it agrees with our experimental observations of collagen content after 12 days in control explants. Therefore, this basal loss would account for collagen degradation that may result from explants culturing rather than injury-driven enzymatic degradation. Accordingly, the free-swelling control cartilage model was modeled with only the constant rate of basal loss *k*_basal_ (see Fig 1d).

Therefore, simulating the net change in collagen content in response to injury, considering: 1) MMP secretion due to cell damage, 2) mitigating MMPs activity by aggrecan[28], and 3) basal *ex vivo* control loss, was given by:

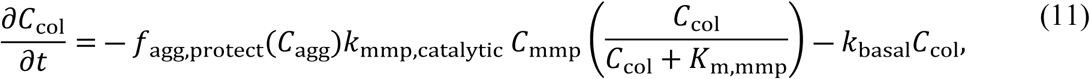

where *k*_mmp,catalytic_defines the rate of MMP catalytic activity, *K*_m,mmp_ represents Michaelis constant for MMPs which describes the MMP concentration through which the MMP–collagen-binding reaction occurs at half-maximum rate[28]. For more information on the parameter values used in the biochemical model, please see the supplementary material (Table S1).

#### 2.4.2 Boundary and initial conditions

For all the variables, zero flux boundary conditions were defined on the bottom surface of the explant, assuming no mass transfer to occur across this surface. For intact aggrecan, degraded aggrecan, and MMPs, Robin boundary conditions were defined through the top and lateral surfaces (representing concentration gradient normal to the surface interface)[28]. For degraded collagen and aggrecanases, Dirichlet boundary conditions were defined at these same surfaces (zero concentration)[28]. The initial healthy cell concentration before the simulation was assumed spatially homogeneous as in immature bovine articular cartilage[28]. The damaged cell concentration was described in Equation (6), and the concentrations for MMPs, aggrecanases, degraded collagen, and degraded aggrecan were assumed to be initially zero.

The initial depth-wise collagen concentration was implemented based on scaling the collagen content absorption unit obtained from the FTIR measurements (Table 1) with the mean collagen concentration 0.2 moles/m^3^ as reported in Kar et al[28]:

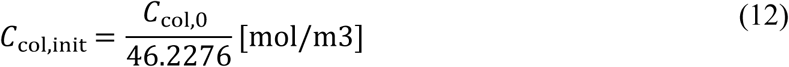

The initial aggrecan distribution was obtained from the fixed charge density (Table 1) as [37]:

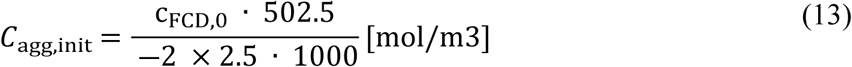

### 2.5 Sensitivity analysis

A sensitivity analysis was conducted for parameters lacking quantitative experimental data but significantly influencing collagen loss in the model. This included parameters governing the extent of cell damage (*k*_INJ_ and *ɛ*_max_), and parameters regulating the enzymatic cleavage of collagen by MMPs (*k*_rate,mmp_, *k*_act_, *k*_mmp_, and *k*_mmp,catalytic_). Reference values for these parameters were selected based on preliminary simulations (Table 2), where the simulated collagen content was matched with the experimentally measured depth-wise collagen content[22].

**Table 2.** **Biochemical model** parameters used for sensitivity analysis. **Bolded values** represent the reference value. Most of the parameters were analysed with ±50% value range. *k*_act_was also tested with 100 times higher value (in red) since no difference was observed within the ±50% range.

| Material Parameter | Range | Description | Reference |
| --- | --- | --- | --- |
| $k_{INJ}$<br>(—) | 0.2, **0.35**, 0.5 | Maximum fraction of healthy cells turning damaged after injury (**Eq 6**) | [7] |
| $\epsilon_{dmg,max}$<br>(%) | 100%, **150%**, 200% | Maximum shear strain threshold for maximum cell damage (**Eq 5**) | [18], model fit |
| $k_{rate,mmp}$<br>($10^{-5}$ 1/s) | 1.8, **3.6**, 5.3 | Rate constant for MMPs generation from damaged cells (**Eq 9**) | model fit, [28] |
| $k_{act}$<br>($10^{-5}$ mol/m<sup>3</sup>) | 1.5, **3**, 4.5, **300** | Aggrecan concentration at half-maximum of MMP activity (**Eq 10**) | [28] |
| $k_{mmp}$<br>($10^{-21}$ mol) | 0.17, **0.35**, 0.52 | Production of MMPs by damaged cells (**Eq 8**) | model fit |
| $k_{mmp,catalytic}$<br>(1/s) | 0.75, **1.5**, 2.25 | Catalytic activity rate of MMPs with collagen fibrils (**Eq 11**) | [28] |

## 3. Results

The control model predicted on average a 19% decrease in the bulk tissue collagen content over 12 days (Fig 2a). On the other hand, in the experiment the bulk tissue collagen content was on average 20% (±6%) smaller on day 12 compared to day 0 (Fig 2b). The depth-wise estimates of collagen content in the control model fitted withing the 95% confidence interval of the experimental results at most of tissue normalized depth (Fig 2c).

**Fig. 2.**
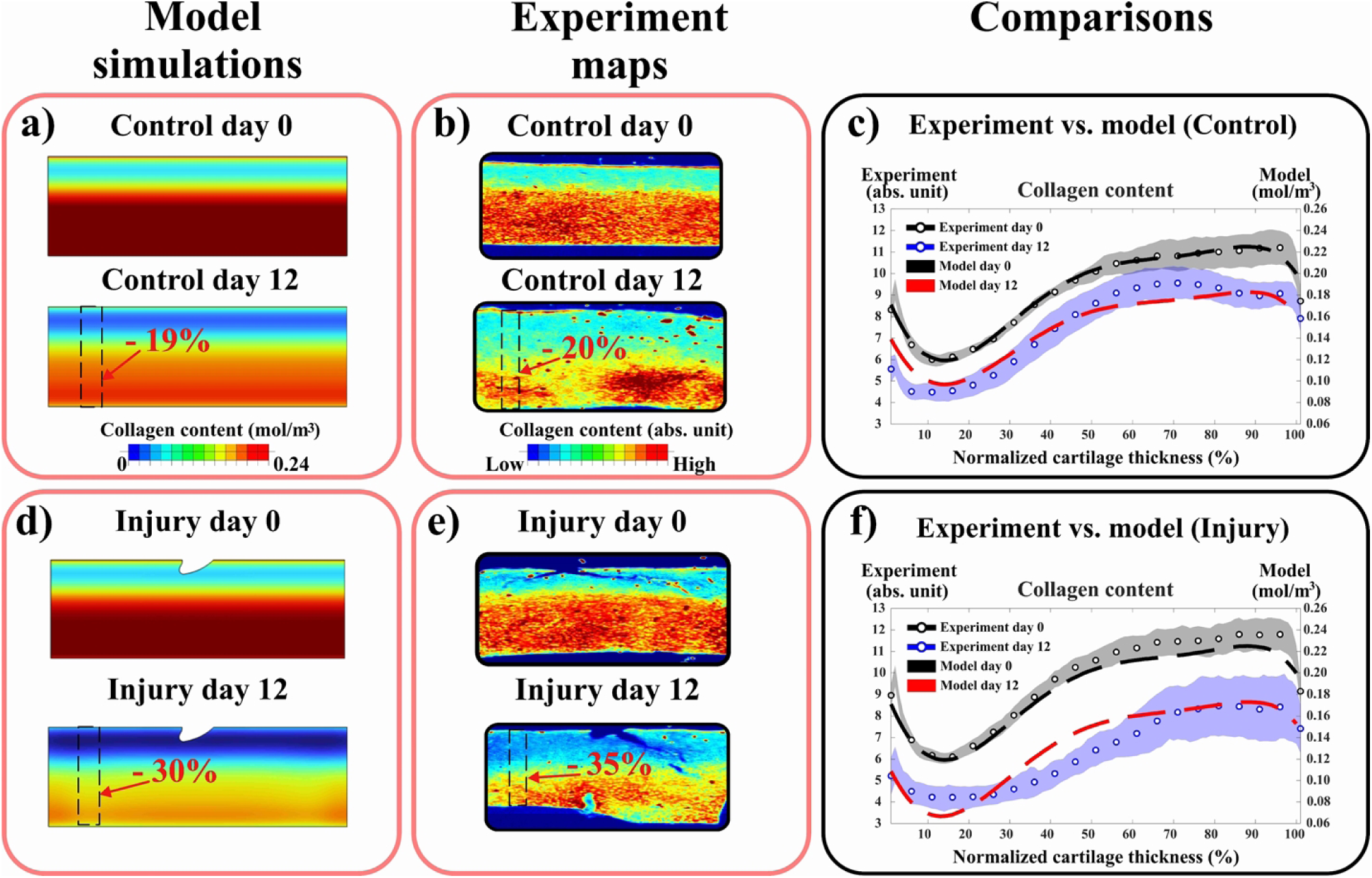
Comparison of experimental and simulated depth-wise collagen content loss. a)&b) collagen distribution overtime in the free-swelling control cartilage in the model and experiment, respectively. c) comparison between model and experimental results along 1 mm-thickness control cartilage showing average 19% bulk decrease in collagen content over 12 days in the model compared to 20% decrease in the experiment. d)&e) collagen distribution overtime in the injuriously loaded cartilage in the model and experiment, respectively. f) the injury model matched with the experimental depth-wise collagen content on day 12, showing on average 30% bulk decrease in collagen content by day 12 compared to 35% decrease in the experiment. The experimental findings are mean ±95% confidence intervals.

The injury model predicted on average 30% (±7%) decrease in bulk collagen content (Fig 2d). On the other hand, the experimental injury group on day 12, compared to day 0, showed an average decrease of 35% (±6%) in collagen content (Fig 2e). In these simulations, depth-wise estimates of collagen content were mostly within the 95% confidence interval of the experimental results (at 0‒30 % and 60‒100% of normalized tissue depth, Fig 2f). Detailed comparisons of the injury model with and without the basal loss term are available in supplementary Material S2 section (Fig S1). Additionally, results for aggrecan loss in experiment and computational modelling were reported elsewhere in our previous studies[36,40].

Sensitivity analysis done on the injury model revealed that increasing or decreasing the shear strain threshold for maximum cell damage (from *ɛ*_dmg,max_ = 150% to *ɛ*_dmg,max_ = 200% or 100%) had a negligible effect on collagen loss (Figure 3a). On the other hand, increasing the maximum allowed cell damage (from *k*_INJ_ = 0.35 to *k*_INJ_ = 0.5) decreased collagen content on day 12 by ∼15% (from 30% to 45% decrease) mostly in the superficial region (at 0‒22% of the normalized tissue depth, Fig 3b).

**Fig. 3.**
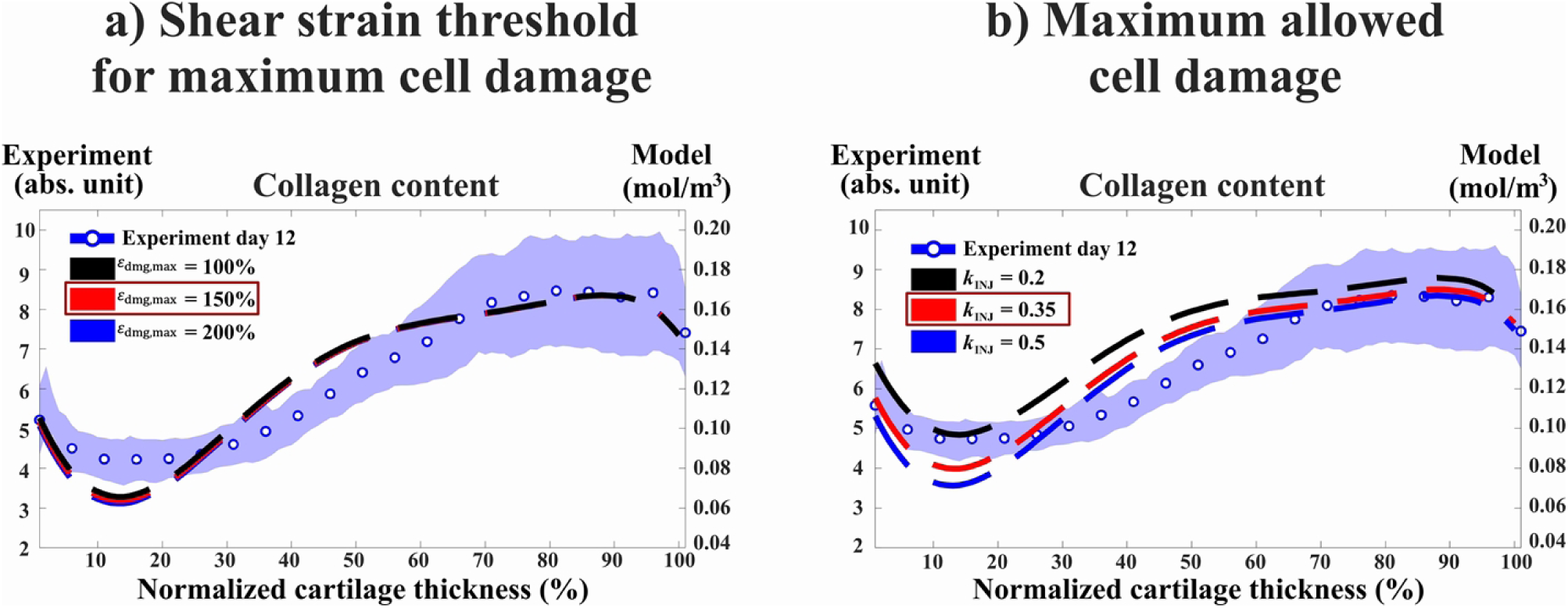
Sensitivity analysis for parameters controlling cell damage in the injury model. a) Increasing or decreasing the shear strain threshold for maximum cell damage by 50% had negligible effect on depth-wise collagen content. b) Increasing the maximum fraction of healthy cells turning damaged (*k*_INJ_) from reference value of 0.35 to 0.5 resulted in increased collagen content loss by ∼15% in the superficial region of cartilage (∼0–25% of normalized depth) while decreasing the value to 0.2 decreased the loss by ∼25% in the superficial cartilage. The red outline highlights the parameter value used in the reference model. The experimental findings are mean ±95% confidence intervals.

Increasing the rate of MMP generation by damaged cells from *k*_rate,mmp_ = 3.6 × 10 ^―5^ 1/s to *k*_rate,mmp_ = 5.3 × 10 ^―5^ 1/s and MMP catalytic activity rate from *k*_mmp,catalytic_ = 1.5 1/s to *k*_mmp,catalytic_ = 2.25 1/s decreased collagen content in both cases on average by 17% (from 30% to 47% decrease), with the most collagen loss occurring at the normalized tissue depth of 0–20% (Fig 4a and Fig 4b). Increasing the amount of MMPs released from damaged cells from *k*_mmp_ = 0.35 × 10^―21^ mol to *k*_mmp_ = 0.52 × 10^―21^ mol decreased collagen content by 18% (from 30% to 48% decrease) at 0–25% of the normalized tissue depth (Fig 4c). On the other hand, increasing the parameter for aggrecan concentration at half-maximum of MMP activity from *k*_act_ = 3 × 10 ^―5^mol/m^3^ to *k*_act_ = 4.5 × 10 ^―5^mol/m^3^ (Equation 10) did not affect the collagen concentration over 12 days (Fig 4d). However, an increase in collagen content by ∼8% at 0‒10% of the normalized tissue depth was observed when the parameter value was increased from *k*_act_ = 3 × 10 ^―5^ mol/m^3^ to *k*_act_ = 3 × 10 ^―3^ mol/m^3^.

**Fig. 4.**
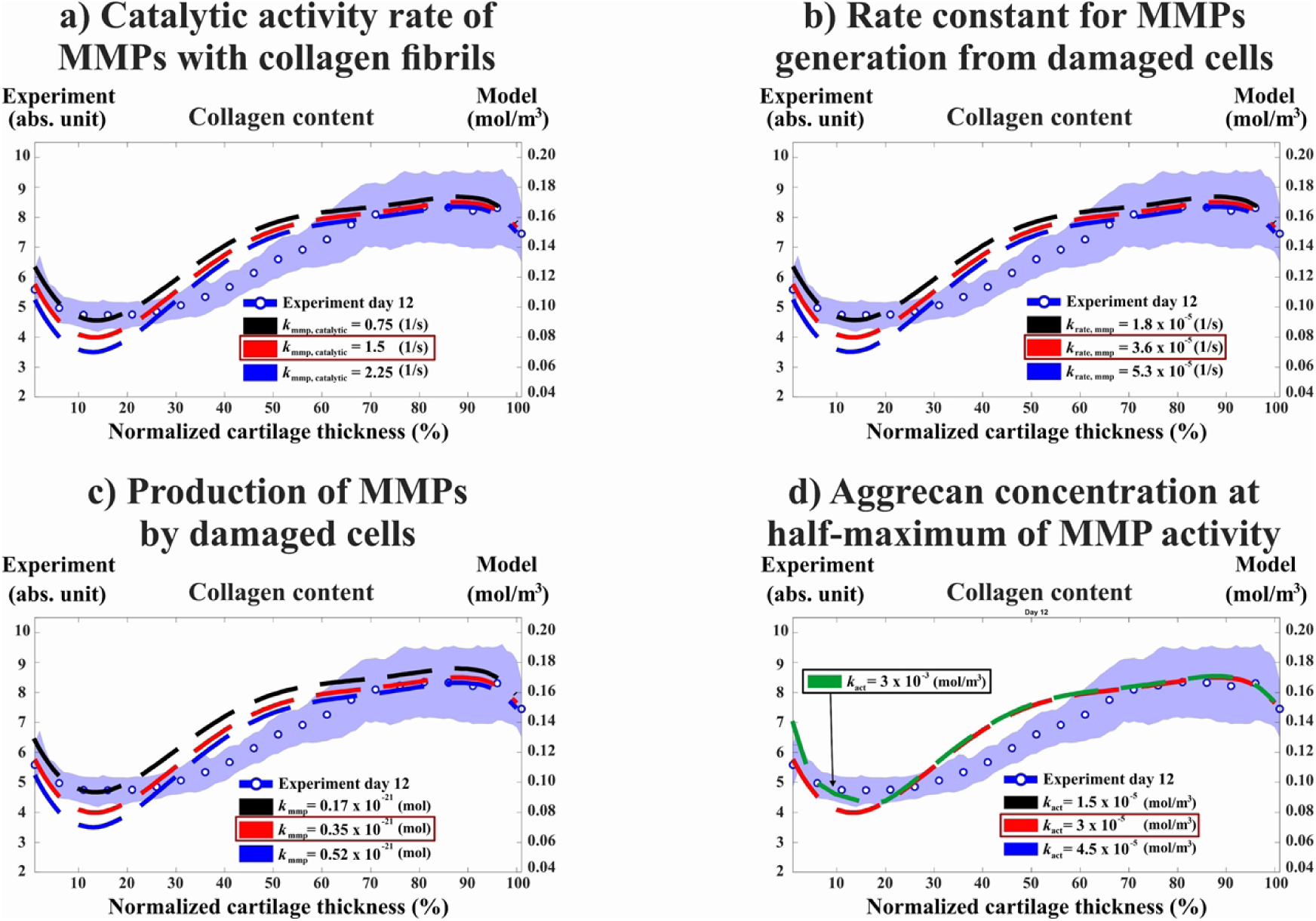
Sensitivity analysis for parameters controlling production and enzymatic activity of MMPs. a)&b) increasing the MMP catalytic rate (*k*_mmp,catalytic_) by 50% and rate constant for MMPs generation from damaged cells (*k*_rate,mmp_) parameters increased the loss of collagen content similarly by ∼17% with the most loss in the superficial region (∼0–20% of normalized depth) as compared to reference model (highlighted in the red outline). c) 50% increased production of MMPs by damaged cells *k*_mmp_ exhibited ∼18% more collagen content loss (∼0 to 25% of normalized depth) compared to reference model. d) changing aggrecan concentration at half-maximum of MMP activity (*k*_act_) parameter had negligible effect on the depth-wise collagen content. However, increasing the value by 100 times more instead of 50% resulted in higher collagen content by ∼8% on average compared to reference model at 0‒10% of normalized tissue depth. The experimental findings are mean ±95% confidence intervals. MMP = matrix metalloproteinase

## 4. Discussion

We created a computational model to simulate cell damage and MMP release caused by injurious loading, and the resulting loss of collagen content in bovine articular cartilage explants over 12 days post-injury. We hypothesized that cell damage in injured cartilage is pronounced in highly strained tissue regions, leading to the activation of MMPs, which subsequently degrade collagen fibrils and result in collagen content loss. The computational injury model predicted depth-wise collagen content loss consistent with experimentally observed data over 12 days (Fig 2)[22]. Our simulations suggest that catabolic cellular activity and the upregulation of MMP activity play a key role in collagen loss during the early stages of cartilage injury.

In this study, no additional mechanical loading was applied to the tissue following the initial injurious loading that would have caused mechanically induced collagen loss or adaptation after the injury. Therefore, the subsequent collagen loss over time was hypothesized to be primarily driven by biochemical factors. The strong agreement between the injury model and the experimental findings supports our proposed mechanism that injury-induced MMP release and enzymatic collagen cleavage is a major contributor to collagen loss[22]. This finding aligns with previous experimental and computational studies investigating biochemically driven degradation in collagen fibril early after excessive and injurious loading[34,45]. Recent computational frameworks have also explored collagen and extracellular matrix loss in cartilage after overloading with attention to the cellular catabolic response, biochemical signaling, and increased proteolytic activity[8,26,34]. For example, Rahman et al.,[34] using 3D FE model, captured significant bulk collagen loss (∼10%) over 3 months due to overloading and MMP-driven degradation. Our model exhibits a faster activation of MMPs and collagen loss, as many of the model parameters were optimized using *ex vivo* experimental data from bovine cartilage, where tissue environment is controlled and the timeframe of cartilage degradation is much faster[1,22]. In contrast, the model by Rahman et al.’s was calibrated using *in vivo* data of cartilage thickness in human subjects and correlating the cartilage thickness to overloading, where cartilage pathological changes would happen in broader timeframe[34]. On the other hand, Kapitanov et al.’s mathematical model addressed the role of blunt impact loading (drop tower impact loading of 2.18 J/cm^2^ energy) on cell damage, the production of reactive oxygen species and the release of interleukin-6 (IL-6) inflammatory cytokines in mature bovine osteochondral explants[33]. Model parameters were calibrated using experimental data on cell viability following the blunt impact loading. Their results showed an increase in IL-6 and reactive oxygen species close to the loaded regions, though the model did not capture matrix loss over a 14-day period. Notably, their model used axial strain to estimate cell damage, which could yield different quantifications of damaged cells. Furthermore, they used homogeneous linear elastic mechanical properties for cartilage with isotropic matrix distribution, which will lead to different mechanical tissue responses compared to our models that incorporated depth-dependent tissue structure.

Our sensitivity analysis revealed that the parameters responsible for the fraction of cell damage (*k*_INJ_) and the production and activity of MMPs (*k*_rate,mmp_, *k*_mmp,catalytic_, and *k*_mmp_) substantially influence the numerical collagen content estimates[13,20,38,46]. However, ±50% variation in *k*_act_ (i.e., aggrecan concentration at half-maximum of MMP activity) yielded no differences in the depth-wise collagen content. We adopted the same parameter value as Kar et al.[28], who demonstrated in their model that a sufficiently high concentration of aggrecan can shield the collagen network from enzymatic degradation caused by IL-1-driven MMP release. However, our study utilized a lower aggrecan concentration (∼half the concentration) compared to their model, potentially limiting the collagen-protecting effect of aggrecan. On the other hand, increasing of *k*_act_ parameter 100 times the reference value resulted in higher collagen content at the superficial zone of the tissue compared to the reference value (Fig 4d). This suggests that further calibration of this parameter is needed. This could enhance our understanding of the mechanistic relationship between aggrecan and collagen in either aggravating or protecting cartilage tissue from the degradation[27]. Also, this could open new avenues for investigating whether potential interventions that target aggrecan biosynthesis in cartilage would play a role in limiting the enzymatic degradation of collagen fibrils [27,36,47].

The predicted collagen content in our injury model deviated from the 95% confidence interval of the experiments at the middle region of the explants (Fig 2b). One reason for this difference might be the experimental setup, where the measurement would not be tracked over time from the same explant (no longitudinal measurements). Although the 0-day and 12-day samples were matched by age and anatomical location, they were not from the exact same patellofemoral groove. As a result, the initial condition used for the 12-day injury model, may not fully match with the actual explants measured at day 12, even though the initial collagen content implemented in the model was obtained from the 0-day control samples of this study.

On the other hand, several degradation mechanisms that we did not consider in the model may also contribute to the depth-dependent loss of collagen content. For instance, previous experimental studies have proposed that mechanical overloading could result in excessive strain in collagen fibrils, potentially leading to rupture of fibrils and subsequent release of collagen fragments to the culture medium[48,49]. Future work could incorporate this hypothesis by including collagen fibril strain into this model framework, combining mechanically and chemically driven factors for collagen loss. Also, our model did not discriminate between the various types of MMPs, each with distinct diffusion coefficients and enzyme kinetics, nor did it explicitly consider spatiotemporal variations in the concentrations of tissue-inhibitors of metalloproteinases. Additionally, we did not discern between different types of cell damage, such as cell apoptosis, mitochondrial dysfunction, or oxidative stress, which could contribute to collagen degradation in distinct temporal and spatial patterns[14,47,50].

The experimental results of the free-swelling control group showed a significant decrease in collagen content on day 12 throughout the entire depth of the tissue compared to day 0 (Fig 2c). Similar observations were reported in previous *ex vivo* studies, where aggrecan loss and cell death were observed in free-swelling cartilage within 12 days[6,36,51,52]. Therefore, this spontaneous basal collagen loss was considered in our workflow with the rate constant *k*_basal_. Including spontaneous collagen loss in the injury model facilitated the differentiation between collagen loss caused in the free-swelling culture conditions and collagen loss caused by the injury-induced MMP release (Fig 2f).

Our model utilizes immature bovine cartilage, which has different material, mechanical, and compositional properties compared to mature human cartilage. Therefore, the experimental findings and model parameter values presented here are not directly translatable to human cartilage. Additionally, the applied loading protocol (strain amplitude, strain rate) can affect the extent of cell damage and the mechano-inflammatory response. Moreover, we assumed constant rates for MMP production and catabolic activity, which may restrict the model’s ability to accurately predict collagen loss at other time intervals[7]. Incorporating new experimental data from other different time points could enhance the calibration of these model parameters, leading to a more accurate simulation of the experiments. We acknowledge that in the context of traumatic joint injuries, damage of cartilage collagen fibrils may also be influenced by inflammatory responses originating from other joint tissues, such as the synovium[24,53,54]. Damage to these tissues can lead to elevated expression of cytokines such as IL-1 and IL-6[24,55], which may subsequently infiltrate cartilage and stimulate MMP production[28]. While our current model focuses on MMP secretion within cartilage in response to direct mechanical injurious loading, this framework can be readily extended to incorporate these exogenous inflammatory pathways. Integrating such mechanisms in future work would enable a more comprehensive simulation of the post-injury cartilage environment.

Lastly, our model did not replicate the previously reported amplified loss of collagen content localized near lesions[22]. This localized collagen degradation was observed to occur rapidly after injury, coinciding with localized cell death on day 0[22,36]. Our model did not incorporate lesion formation and therefore it assumed that lesion is formed directly after the loading. Accordingly, maximum shear strains were not localized at the cartilage surface and were more excessive in deeper regions of the tissue (Fig 1c). However, when considering that lesion is formed during the loading, excessive higher shear strain might be localized near the lesion (see supplementary material, Fig S2). In addition to the release of mechanically ruptured collagen fibrils[22,48], the higher shear strain could lead to higher cell damage and catabolic activity around the lesion compared to intact regions[36], possibly contributing to the amplification of collagen loss near lesions by injury-induced MMP activity [22].

In conclusion, our model underscores the pivotal role of injury-related cell damage and the subsequent downstream MMPs activity to cause the early depth-wise collagen loss observed after injurious loading of cartilage. With additional experimental data and further model calibration and refinement, our model can be valuable for testing the effectiveness of treatment interventions aimed at limiting proteolytic collagen degeneration. This modeling framework could be also implemented into joint-level computational models designed to estimate the progression of PTOA and treatment efficacy[31,56].

## Author contributions

**Moustafa Hamada:** Conceptualization, Data curation, Formal Analysis, Methodology, Visualization, and Writing – Original Draft Preparation. **Atte Eskelinen:** Conceptualization, Methodology, Writing – Review & Editing, and Supervision, Project administration. **Joonas Kosonen:** Conceptualization, Methodology, and Writing – Review & Editing. **Cristina Florea:** Conceptualization, Methodology, and Writing – Review & Editing. **Alan Grodzinsky:** Conceptualization, Methodology, Resources, and Writing – Review & Editing. **Petri Tanska:** Conceptualization, Methodology, Writing – Review & Editing, Supervision, Project administration, Funding acquisition. **Rami Korhonen:** Conceptualization, Methodology, Writing – Review & Editing, Supervision, Project administration, Funding acquisition.

## Data availability statement

All relevant data are within the manuscript and its Supporting information files, and can be accessed via Fairdata research services: https://ida.fairdata.fi/s/NOT-FOR-PUBLICATION-JGKHnzxnFtYH. Upon acceptance of the manuscript for publication, the full set of metadata, codes and models will be made publicly available and the accession numbers/DOI will be provided (Fairdata Research Data Storage Service, https://etsin.fairdata.fi/).

## Ethics statement

The experimental data represented is from bovine animals obtained from a slaughterhouse. Therefore, no ethical permission was needed.

## Supporting information

**Table S1. Biochemical model parameters.**

**Fig. S1. Comparison of the injury model with and without the basal loss of collagen content.**

**Fig. S2. Comparison of the biomechanical model with and without lesion geometry.**

## References

[1] J. Lieberthal, N. Sambamurthy, C.R. Scanzello, Inflammation in joint injury and post-traumatic osteoarthritis, Osteoarthritis Cartilage 23 (2015) 1825–1834. 10.1016/j.joca.2015.08.015.

[2] L.J. Wang, N. Zeng, Z.P. Yan, J.T. Li, G.X. Ni, Post-traumatic osteoarthritis following ACL injury, Arthritis Res Ther 22 (2020). 10.1186/s13075-020-02156-5.

[3] E.M. Roos, Joint injury causes knee osteoarthritis in young adults, (2005).

[4] J.P. Kosonen, A.S.A. Eskelinen, G.A. Orozco, P. Nieminen, D.D. Anderson, A.J. Grodzinsky, R.K. Korhonen, P. Tanska, Injury-related cell death and proteoglycan loss in articular cartilage: Numerical model combining necrosis, reactive oxygen species, and inflammatory cytokines, PLoS Comput Biol 19 (2023). 10.1371/journal.pcbi.1010337.

[5] M.R. Hines, J.E. Goetz, P.C. Gomez-Contreras, S.N. Rodman, S. Liman, E.L. Femino, P.N. Kluz, B.A. Wagner, G.R. Buettner, E.E. Kelley, M.C. Coleman, Extracellular biomolecular free radical formation during injury, Free Radic Biol Med 188 (2022) 175–184. 10.1016/j.freeradbiomed.2022.06.223.

[6] M.A. DiMicco, P. Patwari, P.N. Siparsky, S. Kumar, M.A. Pratta, M.W. Lark, Y.J. Kim, A.J. Grodzinsky, Mechanisms and Kinetics of Glycosaminoglycan Release Following In Vitro Cartilage Injury, Arthritis Rheum 50 (2004) 840–848. 10.1002/art.20101.

[7] J.R. Lyman, J.D. Chappell, T.I. Morales, S.S. Kelley, G.M. Lee, Response of Chondrocytes to Local Mechanical Injury in an Ex Vivo Model, Cartilage 3 (2012) 58–69. 10.1177/1947603511421155.

[8] G.I. Kapitanov, X. Wang, B.P. Ayati, M.J. Brouillette, J.A. Martin, Linking cellular and mechanical processes in articular cartilage lesion formation: A mathematical model, Front Bioeng Biotechnol 4 (2016). 10.3389/fbioe.2016.00080.

[9] P.F. Argote, J.T. Kaplan, A. Poon, X. Xu, L. Cai, N.C. Emery, D.M. Pierce, C.P. Neu, Chondrocyte viability is lost during high-rate impact loading by transfer of amplified strain, but not stress, to pericellular and cellular regions, Osteoarthritis Cartilage 27 (2019) 1822–1830. 10.1016/j.joca.2019.07.018.

[10] E. Brauchle, S. Thude, S.Y. Brucker, K. Schenke-Layland, Cell death stages in single apoptotic and necrotic cells monitored by Raman microspectroscopy, Sci Rep 4 (2014). 10.1038/srep04698.

[11] L. Ramage, G. Nuki, D.M. Salter, Signalling cascades in mechanotransduction: Cell-matrix interactions and mechanical loading, Scand J Med Sci Sports 19 (2009) 457–469. 10.1111/j.1600-0838.2009.00912.x.

[12] T.M. Quinn, A.J. Grodzinsky, E.B. Hunziker, J.D. Sandy, Effects of injurious compression on matrix turnover around individual cells in calf articular cartilage explants, Journal of Orthopaedic Research 16 (1998) 490–499. 10.1002/jor.1100160415.

[13] R.C. Billinghurst, L. Dahlberg, M. Ionescu, A. Reiner, R. Bourne, C. Rorabeck, P. Mitchell, J. Hambor, O. Diekmann, H. Tschesche, J. Chen, H. Van Wart, A.R. Poole, Enhanced Cleavage of Type II Collagen by Collagenases in Osteoarthritic Articular Cartilage, 1997.

[14] M.C. Coleman, P.S. Ramakrishnan, M.J. Brouillette, J.A. Martin, Injurious Loading of Articular Cartilage Compromises Chondrocyte Respiratory Function, Arthritis and Rheumatology 68 (2016) 662–671. 10.1002/art.39460.

[15] J. van Meurs, P. van Lent, A. Holthuysen, D. Lambrou, E. Bayne, I. Singer, W. van den Berg, Active Matrix Metalloproteinases Are Present in Cartilage During Immune Complex-Mediated Arthritis: A Pivotal Role for Stromelysin-1 in Cartilage Destruction, The Journal of Immunology 163 (1999) 5633–5639. 10.4049/jimmunol.163.10.5633.

[16] D.J. Leong, X.I. Gu, Y. Li, J.Y. Lee, D.M. Laudier, R.J. Majeska, M.B. Schaffler, L. Cardoso, H.B. Sun, Matrix metalloproteinase-3 in articular cartilage is upregulated by joint immobilization and suppressed by passive joint motion, Matrix Biology 29 (2010) 420–426. 10.1016/j.matbio.2010.02.004.

[17] H. Bin Sun, R. Nalim, H. Yokota, Expression and activities of matrix metalloproteinases under oscillatory shear in IL-1-stimulated synovial cells, Connect Tissue Res 44 (2003) 42–49. 10.1080/03008200390151954.

[18] S.M. Hosseini, W. Wilson, K. Ito, C.C. Van Donkelaar, A numerical model to study mechanically induced initiation and progression of damage in articular cartilage, Osteoarthritis Cartilage 22 (2014) 95–103. 10.1016/j.joca.2013.10.010.

[19] J. Sanchez-Adams, H.A. Leddy, A.L. McNulty, C.J. O’Conor, F. Guilak, The Mechanobiology of Articular Cartilage: Bearing the Burden of Osteoarthritis, Curr Rheumatol Rep 16 (2014) 1–9. 10.1007/s11926-014-0451-6.

[20] Q. Hu, M. Ecker, Overview of MMP-13 as a promising target for the treatment of osteoarthritis, Int J Mol Sci 22 (2021) 1–22. 10.3390/ijms22041742.

[21] F. Guilak, R.J. Nims, A. Dicks, C.L. Wu, I. Meulenbelt, Osteoarthritis as a disease of the cartilage pericellular matrix, Matrix Biology 71–72 (2018) 40–50. 10.1016/j.matbio.2018.05.008.

[22] M. Hamada, A.S.A. Eskelinen, C. Florea, S. Mikkonen, P. Nieminen, A.J. Grodzinsky, P. Tanska, R.K. Korhonen, Loss of collagen content is localized near cartilage lesions on the day of injurious loading and intensified on day 12, Journal of Orthopaedic Research (2024). 10.1002/jor.25975.

[23] A.M. Loening, I.E. James, M.E. Levenston, A.M. Badger, E.H. Frank, B. Kurz, M.E. Nuttall, H.H. Hung, S.M. Blake, A.J. Grodzinsky, M.W. Lark, Injurious mechanical compression of bovine articular cartilage induces chondrocyte apoptosis, Arch Biochem Biophys 381 (2000) 205–212. 10.1006/abbi.2000.1988.

[24] R.K. Davidson, J.G. Waters, L. Kevorkian, C. Darrah, A. Cooper, S.T. Donell, I.M. Clark, Expression profiling of metalloproteinases and their inhibitors in synovium and cartilage, Arthritis Res Ther 8 (2006). 10.1186/ar2013.

[25] A. Mixon, A. Savage, A.S. Bahar-Moni, M. Adouni, T. Faisal, An in vitro investigation to understand the synergistic role of MMPs-1 and 9 on articular cartilage biomechanical properties, Sci Rep 11 (2021). 10.1038/s41598-021-93744-1.

[26] M. Gierig, P. Wriggers, M. Marino, Computational model of damage-induced growth in soft biological tissues considering the mechanobiology of healing, Biomech Model Mechanobiol 20 (2021) 1297–1315. 10.1007/s10237-021-01445-5.

[27] M.A. Pratta, W. Yao, C. Decicco, M.D. Tortorella, R.Q. Liu, R.A. Copeland, R. Magolda, R.C. Newton, J.M. Trzaskos, E.C. Arner, Aggrecan Protects Cartilage Collagen from Proteolytic Cleavage, Journal of Biological Chemistry 278 (2003) 45539–45545. 10.1074/jbc.M303737200.

[28] S. Kar, D.W. Smith, B.S. Gardiner, Y. Li, Y. Wang, A.J. Grodzinsky, Modeling IL-1 induced degradation of articular cartilage, Arch Biochem Biophys 594 (2016) 37–53. 10.1016/j.abb.2016.02.008.

[29] A.S.A. Eskelinen, P. Tanska, C. Florea, G.A. Orozco, P. Julkunen, A.J. Grodzinsky, R.K. Korhonen, Mechanobiological model for simulation of injured cartilage degradation via proinflammatory cytokines and mechanical, PLoS Comput Biol 16 (2020). 10.1371/journal.pcbi.1007998.

[30] M. Taffetani, M. Griebel, D. Gastaldi, S.M. Klisch, P. Vena, Poroviscoelastic finite element model including continuous fiber distribution for the simulation of nanoindentation tests on articular cartilage, J Mech Behav Biomed Mater 32 (2014) 17–30. 10.1016/j.jmbbm.2013.12.003.

[31] M.E. Mononen, P. Tanska, H. Isaksson, R.K. Korhonen, A novel method to simulate the progression of collagen degeneration of cartilage in the knee: Data from the osteoarthritis initiative, Sci Rep 6 (2016). 10.1038/srep21415.

[32] W. Wilson, C.C. Van Donkelaar, B. Van Rietbergen, K. Ito, R. Huiskes, Stresses in the local collagen network of articular cartilage: A poroviscoelastic fibril-reinforced finite element study, J Biomech 37 (2004) 357–366. 10.1016/S0021-9290(03)00267-7.

[33] G.I. Kapitanov, B.P. Ayati, J.A. Martin, Modeling the effect of blunt impact on mitochondrial function in cartilage: Implications for development of osteoarthritis, PeerJ 2017 (2017). 10.7717/peerj.3468.

[34] M.M. Rahman, P.N. Watton, C.P. Neu, D.M. Pierce, A chemo-mechano-biological modeling framework for cartilage evolving in health, disease, injury, and treatment, Comput Methods Programs Biomed 231 (2023). 10.1016/j.cmpb.2023.107419.

[35] Y. Li, Y. Wang, S. Chubinskaya, B. Schoeberl, E. Florine, P. Kopesky, A.J. Grodzinsky, Effects of insulin-like growth factor-1 and dexamethasone on cytokine-challenged cartilage: Relevance to post-traumatic osteoarthritis, Osteoarthritis Cartilage 23 (2015) 266–274. 10.1016/j.joca.2014.11.006.

[36] ASA. Eskelinen, C. Florea, P. Tanska, HK. Hung, EH. Frank, S. Mikkonen, P. Nieminen, P. Julkunen, AJ. Grodzinsky, RK. Korhonen, Cyclic loading regime considered beneficial does not protect injured and interleukin-1-inflamed cartilage from post-traumatic osteoarthritis, J Biomech (2022) 111181. 10.1016/j.jbiomech.2022.111181.

[37] G.A. Orozco, P. Tanska, C. Florea, A.J. Grodzinsky, R.K. Korhonen, A novel mechanobiological model can predict how physiologically relevant dynamic loading causes proteoglycan loss in mechanically injured articular cartilage, Sci Rep 8 (2018). 10.1038/s41598-018-33759-3.

[38] P.A. West, P.A. Torzilli, C. Chen, P. Lin, N.P. Camacho, Fourier transform infrared imaging spectroscopy analysis of collagenase-induced cartilage degradation, J Biomed Opt 10 (2005) 014015. 10.1117/1.1854131.

[39] X. Bi, X. Yang, M.P.G. Bostrom, N.P. Camacho, Fourier transform infrared imaging spectroscopy investigations in the pathogenesis and repair of cartilage, Biochim Biophys Acta Biomembr 1758 (2006) 934–941. 10.1016/j.bbamem.2006.05.014.

[40] A.S.A. Eskelinen, J.P. Kosonen, M. Hamada, A. Esrafillian, C. Florea, A.J. Grodzinsky, P. Tanska, R.K. Korhonen, Time-dependent computational model of post-traumatic osteoarthritis to estimate how mechanoinflammatory mechanisms impact cartilage aggrecan content, preprint (2024). 10.1101/2024.10.08.617186 (accessed November 27, 2024).

[41] W. Wilson, C.C. Van Donkelaar, B. Van Rietbergen, R. Huiskes, A fibril-reinforced poroviscoelastic swelling model for articular cartilage, J Biomech 38 (2005) 1195–1204. 10.1016/j.jbiomech.2004.07.003.

[42] B. Rolauffs, C. Muehleman, J. Li, B. Kurz, K.E. Kuettner, E. Frank, A.J. Grodzinsky, Vulnerability of the superficial zone of immature articular cartilage to compressive injury, Arthritis Rheum 62 (2010) 3016–3027. 10.1002/art.27610.

[43] E. Han, S.S. Chen, S.M. Klisch, R.L. Sah, Contribution of proteoglycan osmotic swelling pressure to the compressive properties of articular cartilage, Biophys J 101 (2011) 916–924. 10.1016/j.bpj.2011.07.006.

[44] D.D. Dlima, S. Hashimoto, P.C. Chen, C.W. Colwell, M.K. Lotz, Human chondrocyte apoptosis in response to mechanical injury, Osteoarthritis Cartilage 9 (2001) 712–719. 10.1053/joca.2001.0468.

[45] L. Henao-Murillo, K. Ito, C.C. van Donkelaar, Collagen Damage Location in Articular Cartilage Differs if Damage is Caused by Excessive Loading Magnitude or Rate, Ann Biomed Eng 46 (2018) 605–615. 10.1007/s10439-018-1986-x.

[46] G. Jin, R.L. Sah, Y.S. Li, M. Lotz, J.Y.J. Shyy, S. Chien, Biomechanical regulation of matrix metalloproteinase-9 in cultured chondrocytes, Journal of Orthopaedic Research 18 (2000) 899–908. 10.1002/jor.1100180608.

[47] J. Riegger, H. Joos, H.G. Palm, B. Friemert, H. Reichel, A. Ignatius, R.E. Brenner, Antioxidative therapy in an ex vivo human cartilage trauma-model: attenuation of trauma-induced cell loss and ECM-destructive enzymes by N-acetyl cysteine, Osteoarthritis Cartilage 24 (2016) 2171–2180. 10.1016/j.joca.2016.07.019.

[48] M. Thibault ’, A.R. Poole ’, M.D. Buschmann, Cyclic compression of cartilagdbone explants in vitro leads to physical weakening, mechanical breakdown of collagen and release of matrix fragments, 2002. www.elsevier.com/locate/orthres.

[49] C.T. Chen, C.-T., Burton-Wurster, N., Lust, G., Bank, R. A., & Tekoppele, J. M. (1999). Compositional and metabolic changes in damaged cartilage are peak-stress, stress-rate, and loading-duration dependent. Journal of Orthopaedic Research, 17(6), 870–879. 10.1002/jor.1100170612.

[50] M.J. Brouillette, P.S. Ramakrishnan, V.M. Wagner, E.E. Sauter, B.J. Journot, T.O. McKinley, J.A. Martin, Strain-dependent oxidant release in articular cartilage originates from mitochondria, Biomech Model Mechanobiol 13 (2014) 565–572. 10.1007/s10237-013-0518-8.

[51] P. Patwari, M.N. Cook, M.A. DiMicco, S.M. Blake, I.E. James, S. Kumar, A.A. Cole, M.W. Lark, A.J. Grodzinsky, Proteoglycan degradation after injurious compression of bovine and human articular cartilage in vitro: Interaction with exogenous cytokines, Arthritis Rheum 48 (2003) 1292–1301. 10.1002/art.10892.

[52] B. Kurz, M. Jin, P. Patwari, D.M. Cheng ’, M.W. Lark ’, A.J. Grodzinsky, Biosynthetic response and mechanical properties of articular cartilage after injurious compression, 2001. www.elsevier.com/locate/orthres.

[53] Y. Sui, J.H. Lee, M.A. DiMicco, E.J. Vanderploeg, S.M. Blake, H.H. Hung, A.H.K. Plaas, I.E. James, X.Y. Song, M.W. Lark, A.J. Grodzinsky, Mechanical injury potentiates proteoglycan catabolism induced by interleukin-6 with soluble interleukin-6 receptor and tumor necrosis factor α in immature bovine and adult human articular cartilage, Arthritis Rheum 60 (2009) 2985–2996. 10.1002/art.24857.

[54] J.M. Dayer, The pivotal role of interleukin-1 in the clinical manifestations of rheumatoid arthritis, Rheumatology 42 (2003). 10.1093/rheumatology/keg326.

[55] M.L. Redondo, D.R. Christian, A.B. Yanke, The Role of Synovium and Synovial Fluid in Joint Hemostasis, in: Joint Preservation of the Knee, Springer International Publishing, Cham, 2019: pp. 57–67. 10.1007/978-3-030-01491-9_4.

[56] R.K. Korhonen, A.S.A. Eskelinen, G.A. Orozco, A. Esrafilian, C. Florea, P. Tanska, Multiscale In Silico Modeling of Cartilage Injuries, in: Adv Exp Med Biol, Springer, 2023: pp. 45–56. 10.1007/978-3-031-25588-5_3.

[57] L. Li, M.D. Buschmann A. Shirazi-Adl, A fibril reinforced nonhomogeneous poroelastic model for articular cartilage: inhomogeneous response in unconfined compression, Journal of Biomechanics, 2000: 1533–1541.

